# High Calcium Transport by Polycystin-2 (TRPP2) Induces Channel Clustering and Coordinated Oscillatory Behavior

**DOI:** 10.1101/2022.03.02.482677

**Authors:** Irina F. Velázquez, Horacio F. Cantiello, María del Rocío Cantero

## Abstract

The regulation by Ca^2+^ of Ca^2+^-permeable ion channels represents an important mechanism in the control of cell function. Polycystin-2 (PC2, TRPP2), a member of the TRP channel family (*Transient Potential Receptor*), is a Ca^2+^ permeable non-selective cation channel. Previous studies from our laboratory demonstrated that physiological concentrations of Ca^2+^ do not regulate *in vitro* translated PC2 (PC2_iv_) channel activity. However, the issue as to PC2’s Ca^2+^ permeability and regulation remain ill-defined. In this study, we assessed Ca^2+^ transport by PC2_iv,_ in a lipid bilayer reconstitution system in the presence of a high Ca^2+^ gradient (CaCl_2_ 100 mM *cis*, CaCl_2_ 10 mM *trans*). PC2_iv_ channel reconstitution was conducted in the presence of either 3:7 or 7:3 1-palmitoyl-2-oleoyl-choline (POPC) and ethanolamine (POPE) lipid mixtures. Reconstituted PC2_iv_ showed spontaneous Ca^2+^ currents, in both lipid mixtures with a maximum conductance of 63 ± 13 pS (*n* = 19) and 105 pS ± 9.8 (*n* = 9), respectively. In both cases, experimental data were best fitted with the Goldman-Hodgkin-Katz equation, showing a reversal potential (V_rev_ ~ −27 mV) consistent with strict Ca^2+^ selectivity. The R742X mutated PC2 (PC2_R742X_), lacking the carboxy terminal domain of the channel showed no differences with wild type PC2. Interestingly, spontaneous Ca^2+^ current oscillations were observed whenever PC2-containing samples were reconstituted in the 3:7, but not 7:3 POPC:POPE lipid mixture. The amplitude and frequency of the oscillations were highly dependent on the applied voltage, the imposed Ca^2+^ gradient, and the presence of high Ca^2+^, which induced PC2 channel clustering as observed by atomic force microscopy (AFM). We also used the QuB suite to kinetically model the PC2 channel Ca^2+^ oscillations based on the presence of subconductance states in the channel. The encompassed data provide new evidence to support a high Ca^2+^ permeability by PC2, and a novel regulatory feedback mechanism dependent on the presence of Ca^2+^ and phospholipids on its function.

**Statement of Significance:** The regulation by Ca^2+^ of Ca^2+^-permeable ion channels represents an important mechanism in the control of cell function. The *Transient Potential Receptor* channel Polycystin-2 (TRPP2, PC2), is a Ca^2+^ permeable non-selective cation channel. Ca^2+^ transport by PC2 has largely been inferred by changes in reversal potential. This study provides experimental evidence on the Ca^2+^-transporting capabilities of PC2 in high Ca^2+^ that is modulated by lipids and generates a novel phenomenon of oscillatory currents by channel clustering and multiple subconductance behavior. PC2 can be self-regulated by feedback mechanisms, which are independent of external regulatory proteins. This oscillatory behavior, previously unknown for a single channel species, depend on the presence of Ca^2+^ interaction sites as have been postulated for the channel protein.

## Introduction

Transient receptor potential (TRP) channels (1) are considered of fundamental importance for their contribution to the resting potential of cells, and a variety of cellular mechanisms in both excitable and non-excitable cells (2, 3). PC2 is a structural tetramer, whose activity is the summation of subconductance levels corresponding to a sequential activation of each of the four monomers (4, 5). Hetero-oligomerization such as that observed in TRPC1-PC2 hetero-complexes show distinct functional properties, including resistance to low pH, and amiloride sensitivity, not observed in the homo-tetramers (5). PC2 is involved in the entry of Ca^2+^ into different epithelial tissues and organs (4, 6–9). The contribution of PC2 to cell signaling and Ca^2+^ homeostasis has been reported in different cellular compartments, including the primary cilium, intracellular Ca^2+^ reservoirs such as the endoplasmic reticulum (ER), and the plasma membrane (7, 10, 11). Thus, PC2 plays an important role in Ca^2+^-mediated cell activation, and the dynamics of intracellular Ca^2+^ usually shows a complex space-time behavior. Many cell types respond to agonist stimulation with fluctuations in Ca^2+^ concentrations (12), which result from the periodic release of Ca^2+^ stored in intracellular compartments. In many cases, the oscillations are a signal of coded frequency that allows the cell to use Ca^2+^ as a second messenger. A known model of cytoplasmic Ca^2+^ oscillations is the two-reservoir model (“Two Pool”, 13), based on the production of oscillations through the interaction between two Ca^2+^ reservoirs, one sensitive, and another insensitive to IP3 (sensitive to Ca^2+^). While a stimulus is present, a constant influx of Ca^2+^ from the insensitive IP3 reservoir occurs (14). Considering the fact that PC2 is present in the plasma membrane and its function is controlled by external Ca^2+^ concentrations (15), it is of interest to take a closer look at the Ca^2+^ permeability of this cation channel.

To date, Ca^2+^ transport by PC2 and homologues has been inferred by either change in reversal potential, and/or changes in ionic conductance at very high Ca^2+^ concentrations (4, 16). An intrinsic PC2 Ca^2+^ permeability at physiological Ca^2+^ was calculated for reconstituted human syncytiotrophoblast PC2 (PC2_hst_) (16). In that study, a reduction of *cis* Ca^2+^ from 10 μM to sub-nanomolar concentrations by addition of either EGTA or BAPTA, decreased the PC2-mediated K^+^ currents. However, similar experiments conducted with *in vitro* translated PC2 (PC2_iv_) showed no changes in channel activity, suggesting that a Ca^2+^ microdomain surrounding the PC2_hst_ regulated the channel. This microdomain, which is absent in the *in vitro* translated channel protein, turned out to be comprised of Ca^2+^-sensitive cytoskeletal proteins that interact with the channel, and confer the regulation by Ca^2+^ (17). Ehrlich’s group reported Ca^2+^-dependent conformational changes in the isolated carboxy terminus of PC2 (18), supporting the idea that the PC2 channel functionally interacts with Ca^2+^. It is currently unknown as to whether this phenomenon transmits any functional property to the channel complex. Our previous data were consistent with a scenario in which Ca^2+^ transport through PC2 was fed back by accessing intracellular cytoplasmic sites, thus maintaining channel function (16). Access to this information and the delivery of physiological concentrations of extracellular Ca^2+^ to the cytoplasmic domain allowed us to quantify the boundaries of Ca^2+^ permeability by PC2 from the relationship between the time necessary to reach PC2’s channel half maximal activity (*t_1/2_*) and the external concentration of Ca^2+^. The flux coefficient (*τ*_Ca_) calculated in KCl and CaCl_2_ between 0.3 nM and 1 mM, was 20 μs, reflecting the average time taken by Ca^2+^ to traverse the channel pore (*trans/cis*) to access and activate the Ca^2+^ regulatory sites. From the estimated *τ*_Ca_ the calculated Ca^2+^ flux (*J*_Ca_) through PC2_hst_ at physiologically relevant Ca^2+^ concentrations were in the order of 9.32 × 10^−20^ moles/s to 1.48 × 10^−18^ moles/s, consistent with a PC2_hst_ Ca^2+^ conductance from ~ 0.15 to 2.38 pS in the presence of 1 mM external Ca^2+^ (16). These flux values are high compared to those reported for most Ca^2+^-permeable channels under physiologically imposed electrochemical conditions (19). Our data were validated by means of electrodiffusional experiments obtained with PC2_hst_ in the presence of a K^+^ gradient and a high *trans* Ca^2+^ concentration (90 mM) (16). We require direct measurements of electrodiffusional Ca^2+^ fluxes through PC2 to confirm these values.

Thus, in the present study we explored the Ca^2+^ permeability of PC2 in the presence of a Ca^2+^ chemical gradient and the absence of monovalent cations. We observed Ca^2+^ transport through PC2 in the isolated protein (PC2_iv_). Interestingly, the PC2_iv_ maximal single channel conductance depended on the lipid mixture composition, showing as much as twice the channel permeability on the ratio of phospholipids. Another interesting finding was the presence of a novel Ca^2+^-induced functional clustering of PC2 channels, observed by AFM, to synchronize Ca^2+^ waves in the presence of a 3:7, but not 7:3 PC:PE lipid mixture. This is to our knowledge the first demonstration of Ca^2+^ oscillations in the presence of a single ion channel species. We constructed a theoretical four-state kinetic model of the PC2-mediated Ca^2+^ oscillations by with the QuB Software suite (20) that depended exclusively on the presence of subconductance states of the channel. The present study provides evidence on a novel regulatory mechanism of Ca^2+^ transport by PC2.

## Methods

### Preparation of in vitro translated PC2

*In vitro* translated PC2 (PC2_iv_) was prepared as previously reported (4). Briefly, the plasmid pGEM-PKD2 encoding PC2, was *in vitro* transcribed and translated with a reticulocyte lysate system TnT T7 (Promega, Fitchburg, WI) by incubation of plasmid DNA (1 mg) and 50 mL of the reaction mixture for 90 min at 30ºC. The PC2_iv_ was introduced by dialysis into liposomes formed by a mixture of 1-palmitoyl-2-oleoyl-choline (POPC): 1-palmitoyl-2-oleoyl-ethanolamine (POPE) in a 7:3 ratio (4, 5). R742X truncated PC2 (PC2_R742X_) lacking the carboxy terminal tail of the channel was prepared as previously described (21).

### Preparation of proteoliposomes

PC2 was introduced into liposomes formed with a mixture of POPC:POPE, 7:3 for its subsequent reconstitution in lipid bilayers. Proteoliposomes containing PC2 were prepared with a 7:3 mixture of the polar lipids POPC (10 mg/ml, Avanti, Inc., AL) and 1.2 ml POPE (25 mg/ml, Avanti, Inc., AL) to obtain a mixture of lipids (40 mg) which was then dried with N_2_ and re-dissolved in n-decane. To the lipid mixture were added 4 ml of cholate buffer (25 mM Na^+^-cholate, 150 mM NaCl, 0.1 mM EDTA, 20 mM HEPES, pH 7.2), which were sonicated in a bath for 45 min. For the preparation of the proteoliposomes the PC2 (1 mg / ml) was diluted to 1: 1.000, and 1 ml was added to 100 μl of the lipid solution described above, and mixed with 0.2 ml of Na^+^-cholate. The mixture was dialyzed against dialysis buffer (150 mM NaCl, 0.1 mM EDTA, 20 mM HEPES, pH 7.2) for 3 days at 4°C (with three buffer changes) (5).

### Ion channel reconstitution

PC2 containing vesicles were inserted into a lipid bilayer reconstitution system as previously reported (4). For the present studies we used a CaCl_2_ chemical gradient (*cis/trans*100:10 mM). Lipid mixtures were prepared from either 3:7 or 7:3 ratios, of synthetic POPC (10 mg/ml) and POPE (10 mg/ml) stock solutions (Avanti Polar Lipids, Birminham, AL), respectively. Lipid mixtures were dried with N_2_ and re-dissolved in 12 μl n-decane, as described (22). Unless otherwise stated, the *cis* chamber contained a solution of CaCl_2_ 100 mM, and HEPES 10 mM, at pH 7.40, while the *trans* side contained a similar solution with lower CaCl_2_ (10 mM), to create a CaCl_2_ chemical gradient. Wherever indicated, PC2 preparations were also reconstituted in the presence of either a biionic gradient K^+^/Ca^2+^ (KCl 150 *cis* and 10 *trans*) or symmetrical K^+^ solution (KCl 150, CaCl_2_ 10 μM *cis* and KCl 150, CaCl_2_ 10 μM *trans*).

### PC2 channel inhibition

PC2_iv_ function was identified at the end of the experiment by inhibition with either *trans* (external) amiloride (100 μM), or *cis* (cytoplasmic side) anti-PC2 antibody (1:1000, 10 μl, sc-25749, Santa Cruz Biotechnology), properties that also ensured its orientation in the reconstituted membrane.

### Data acquisition and analysis

Single channel currents were obtained with a 10 GΩ feedback resistor PC501A patch-clamp amplifier (Warner Instruments, Hamden, CT), which was driven with the software pCLAMP 6.2 (4) from a personal computer. Output (voltage) signals were low-pass-filtered at 700 Hz (3 dB) with an eight-pole, Bessel-type filter (Frequency Devices, Haverhill, MA). Single channel currents were further filtered for display purposes only. Unless otherwise stated, pClamp 10.0 (Axon Instruments, Foster City, CA), and Sigmaplot 11.0 (Jandel Scientific, Corte Madera, CA), were used for data analysis, and statistical analysis and graphics, respectively.

### Atomic force microscopy

PC2 channel complexes were imaged with an ICON AFM attached to a Nano-Scope V controller (Bruker, Sta. Barbara, CA), as previously reported (5, 23). Unless otherwise stated, samples were scanned with oxide sharpened silicon-nitride tips (DNP-S, Bruker). BLMs were scanned in either tapping or contact mode, in liquid (Fig 2) with silicon nitride cantilevers with a spring constant of 0.06 N/m and operating frequencies for the tapping mode of 8 kHz (tip radius estimated as ~25 nm). Scanning rates varied from 0.3 to 2 Hz.

**Fig. 1:**
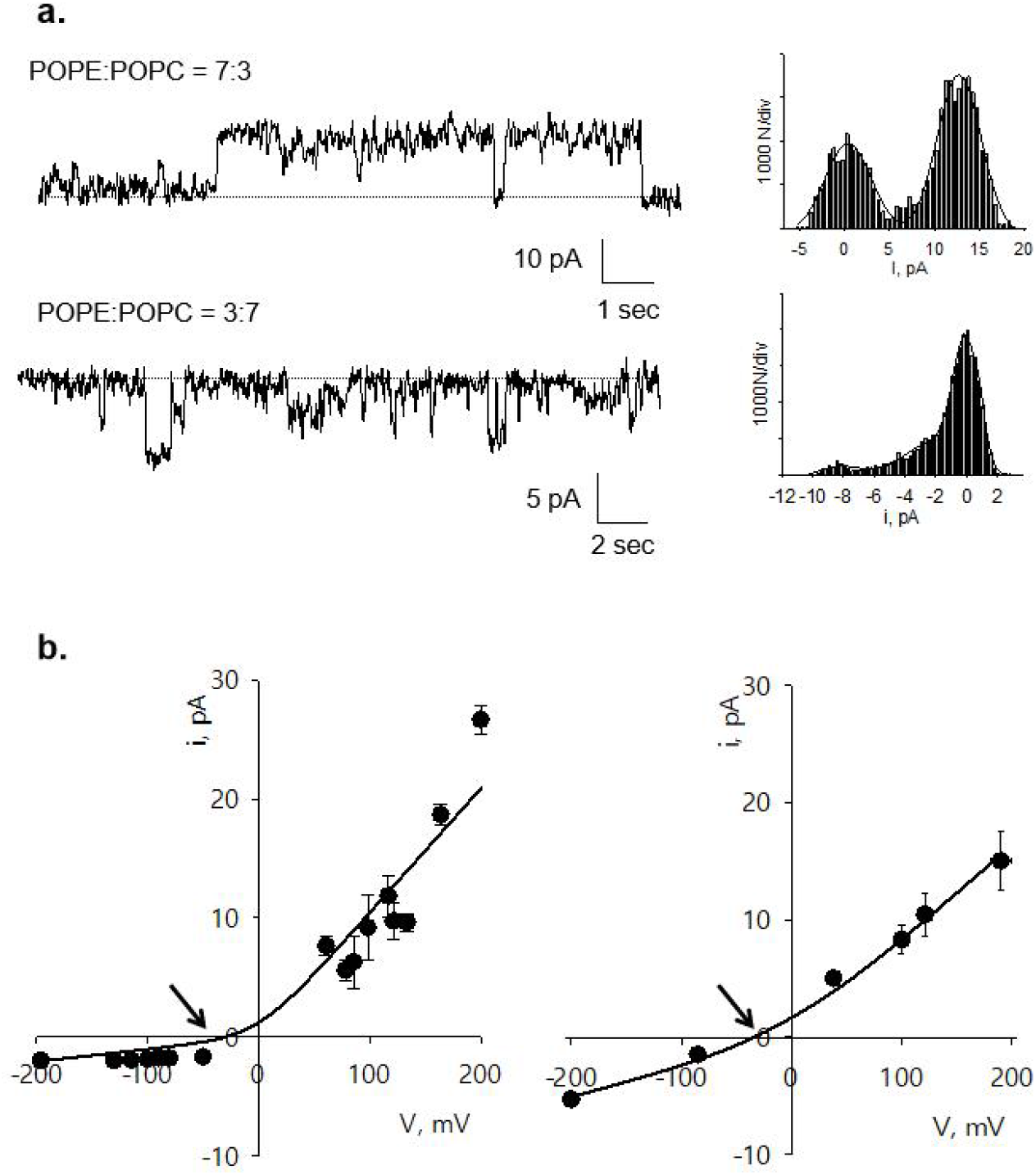
Ca^2+^ transport through PC2_iv_. **a.** Left. Representative PC2_iv_ single channel Ca^2+^ currents. The channel was reconstituted in a lipid bilayer system in the presence of a Ca^2+^ gradient and either a POPE:POPC 7:3 (100 mV) or 3:7 (−200 mV) lipid mixture. Representative tracings from n = 105/360. The closed level is indicated by the dotted line. Right. All-point amplitude histogram. Black line represents the best fit for two Gaussians, corresponding to the closed and open levels. **b.** Current to voltage relationships obtained with either 7:3 (Left) or 3:7 (Right) POPC: POPE lipid mixtures fitted with the GHK model (solid line). Gray symbols represents current-to-voltage relationship for PC2_R742X_.

**Fig. 2:**
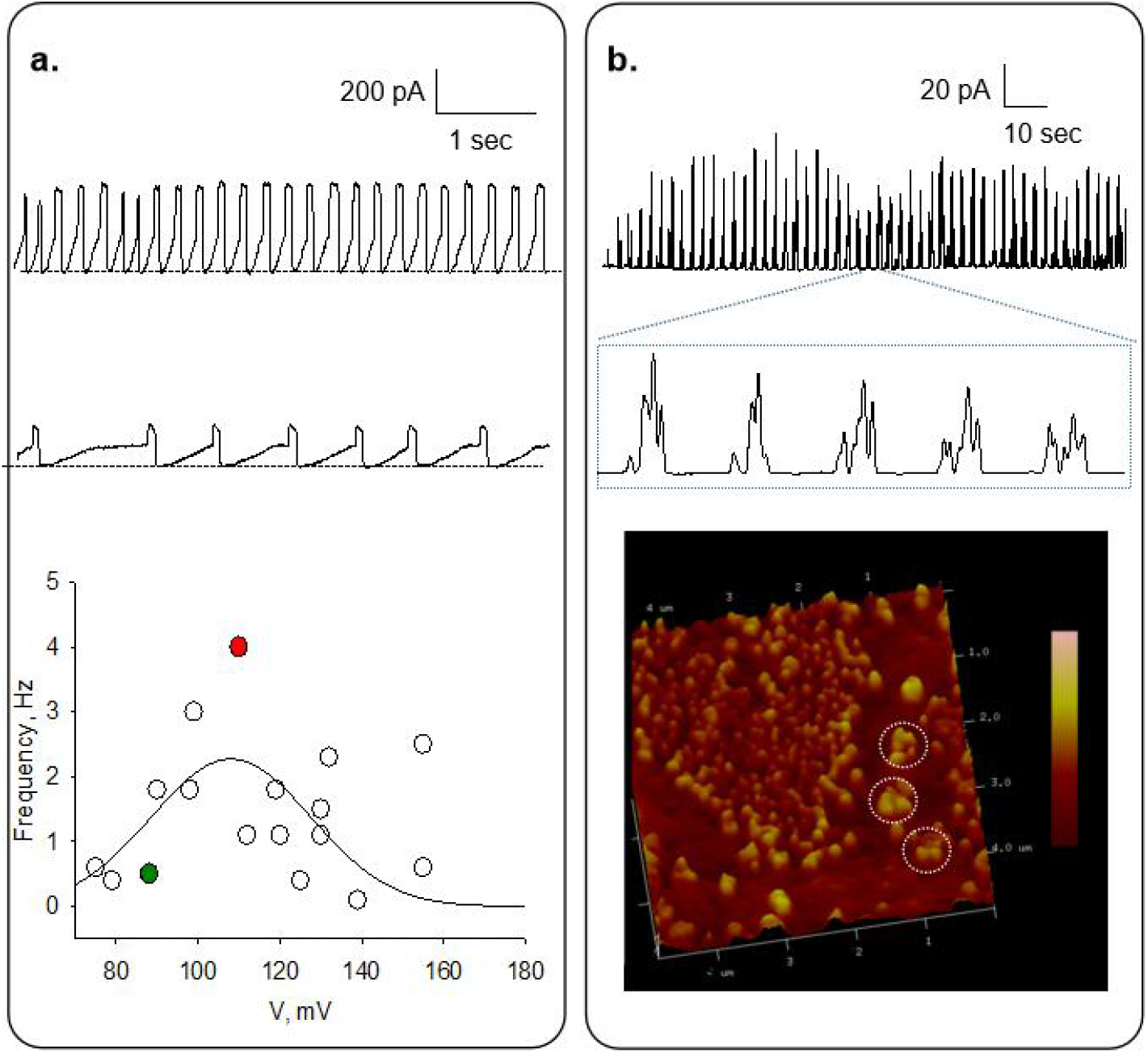
PC2_iv_ Ca^2+^ current oscillations in a CaCl_2_ gradient. **a.** Top and Middle. Representative electric oscillations in the presence of CaCl_2_ 100 mM cis and CaCl_2_ 10 mM trans reconstituted in the presence of the 3:7 POPC: POPE lipid mixture. Zero current level is indicated by the dashed line. Bottom. Frequency of oscillations (Hz) vs. voltage (mV). Data were adjusted with the phenomenological equation (Eq. 3) (black curve). Red and green dots are the values obtained from the Top an Middle tracings. **b.** Electrical recording obtained with the AFM connected to a lipid bilayer system (23). Scanning shows the PC2 channel clustering in high Ca^2+^.

### Ca^2+^ gradients and biionic conditions

Whenever indicated, addition of Ca^2+^ was performed by replacement of 200 μl of solution of the *trans* hemi-chamber by 200 μl CaCl_2_ 1.0 M. Final concentration obtained in the *trans* compartment was 210 mM in the bi-ionic gradient (150 mM KCl *cis*, CaCl_2_ 10 mM *trans*) and 200 mM in symmetrical K^+^ (150 mM KCl *cis*, KCl 150 mM *trans*).

### Kinetic Modeling of the PC2 Channel

Kinetic modeling of PC2 behavior was simulated with the QuB Classic software 2.0.0.13 (State University of New York, Buffalo, NY) (20). Kinetic models of the PC2 channel were simulated and exported to Clampfit 10.5 (Molecular Devices, LLC) for further analysis.

### Goldman-Hodgkin-Katz and other field equations

The ionic conductance of the PC2 channel, was first determined by fitting experimental data from current to voltage relationships to the corrected Goldman-Hodgkin-Katz (GHK) current equation (24) such that:

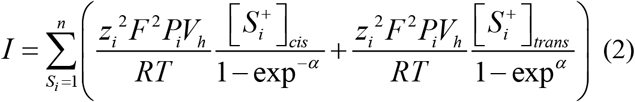

where *I* is the single channel current, [*S_i_^+^*] is the concentration of ion *i* in either (*cis, trans*) chamber, *V*_*h*_ is the holding potential, and α = *z_i_FV_h_/RT*. The constants *F*, *R* and *T* have their usual meaning. The original GHK approximation assumes a linear, or symmetric electric field profile (25, 26) across the membrane (constant field approximation), which is not always supported by channel behavior. In fact, early work by Hodgkin & Keynes (27, 28) observed that the Ussing flux ratio, which also applies to unidirectional current ratios *I*_*i*_/*I*_*o*_ = exp (α) (28, 29) can sometimes present ratios where the exponent containing the electric potential term is multiplied by factors higher than 1. These higher ratios have been associated with multi-occupancy in the channel pore (26), where the number multiplying the exponential term represents the number of ions in the pore, and possibly other explanations such as electrical interactions in the pore, either between ions, or between an ionic species and the walls of the pore.

### Statistical methods

Maximal Ca^2+^ conductance (*g*_*max*_) for PC2_iv_ with either 3:7 or 7:3 POPC:POPE lipid mixture was obtained by linear regression, and comparisons were evaluated by Student t test, with a significance of p < 0.05.

## Results

### Ca^2+^ transport through PC2_iv_

To study Ca^2+^ transport through PC2_iv_, the channel was reconstituted in a lipid bilayer system (see Materials & Methods) in the presence of a CaCl_2_ chemical gradient (100 mM *cis* vs. 10 mM *trans*), in either 3:7 or 7:3 POPC:POPE lipid mixtures. Holding potential differences between ±200 mV were applied. We observed spontaneous single channel Ca^2+^ currents (n = 105/360) in both lipid mixtures (Fig. 1a). Spontaneous single channel currents were most often elicited at high holding potentials (>100 mV) and most frequently, several channel openings were observed at the same time, saturating the amplifier. Usually, we required application of holding potentials as high as 200 mV to elicit ion channel transitions that could be recorded at lower holding potentials afterwards. Current-to-voltage relationships for each lipid mixture revealed a *V*_*rev*_ expected for Ca^2+^ permeation (Theoretical *V*^*Ca*^_*rev*_ of −29.5 mV) (Fig 1b). Data were fitted with the corrected GHK equation (Eq. 2) in the presence of either, 7:3 POPC: POPE, (Fig. 1b Left), or 3:7 (Fig. 1b Right) lipid compositions. In the presence of the 7:3 POPC: POPE mixture, the maximum Ca^2+^ conductance (*g*_*max*_) was 104.4 ± 9.8 pS (*n* = 9), with a *V*_*rev*_ was −28 mV, most consistent with a Ca^2+^ permeability (*P*_*Ca*_) of 6.94 × 10^−14^ cm^3^s^−1^. In the presence of the 3:7 POPC: POPE lipid mixture, however, *g*_*max*_ was 78.8 ± 7.1 pS (*n* = 19), with a *P*_*Ca*_ of 1.18 × 10^−14^ cm^3^s^−1^, thus 83% lower than that with the other lipid mixture (p < 0.05), and a *V*_*rev*_ of −31 mV.

### Ca^2+^ transport through truncated PC2_R742X_

Ca^2+^ transport was also studied through the ADPKD-causing truncated PC2_R742X_. This mutation lacks the cytoplasmic carboxy terminal tail of PC2. PC2_R742X_ was reconstituted using the 3:7 POPC:POPE lipid mixture in the presence of a Ca^2+^ gradient similar to that applied to wild type PC2_iv_. Spontaneous Ca^2+^ currents were observed (*n* = 30/41), with a similar Ca^2+^ conductance to that of the wild type PC2_iv_ (Fig. 1b Right, gray symbols). It is important to note that was very difficult to obtain single channel tracings from PC2_R742X_ reconstituted in Ca^2+^ chemical gradient.

### Current oscillations of reconstituted PC2_iv_ in the presence of a Ca^2+^ gradient

Most often several channel openings were simultaneously observed after reconstitution of PC2_iv_ in a Ca^2+^ gradient (*cis* 100 mM vs. *trans* 10 mM). Under these conditions, an interesting phenomenon was the observation of current oscillations (Fig. 2 a&b). This electrical phenomenon was present in experiments where PC2_iv_ was reconstituted in the 3:7 POPC:POPE lipid mixture (*n* = 29/139). Reconstitution of PC2_iv_ in the 7:3 POPC:POPE lipid mixture, showed single channel activity in 16/96 experiments with no current oscillations. Also interesting, reconstitution of PC2_R472X_ under similar conditions, showed no current oscillations in the 3:7 POPC:POPE lipidic mixture (*n* = 0/41).

The oscillatory current amplitude (in pA) depended on the applied voltage (Fig. 3a Top), being greater at higher potentials. The oscillation frequency (Hz) did not respond linearly with respect to the holding potential (Fig. 2a Bottom), but instead according to the phenomenological equation,

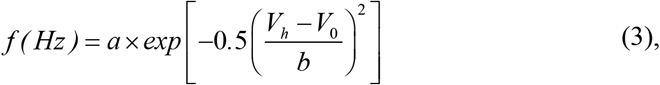

where *f* is the frequency in Hz, *a* and *b* are constants, *V*_*h*_ is the holding potential (in mV) and *V*_*0*_ is the peak voltage (also in mV).

**Fig. 3:**
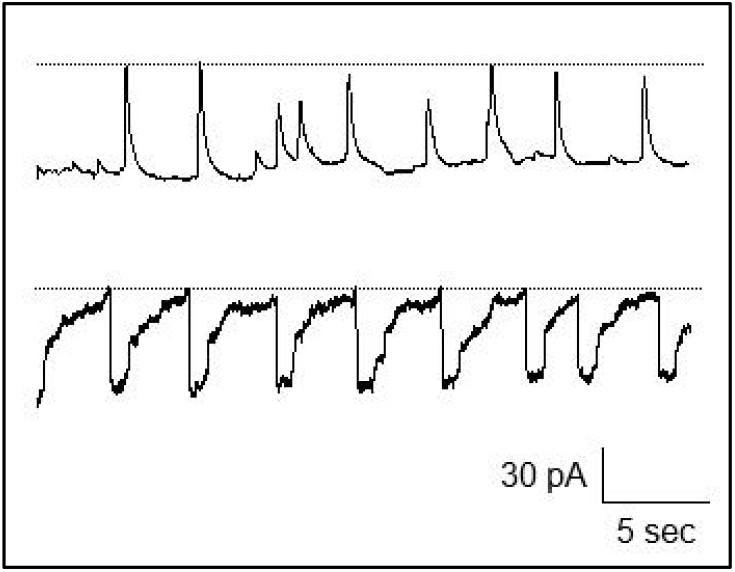
PC2_iv_ current oscillations under several ionic conditions. Representative tracings of PC2_iv_ electric oscillations in the presence of either KCl 150 mM, 10 μM CaCl_2_ cis, 210 mM trans CaCl_2_ (Top) or KCl 150 mM, 10 μM CaCl_2_ cis; 120 mM KCl, CaCl_2_ 200 mM trans (Bottom) reconstituted in a 3:7 POPC:POPE lipid mixture. Zero current level is indicated by the dotted line.

### PC2 channel clustering and Ca^2+^ oscillations

To further explore whether the lipid composition could be implicated in generating the Ca^2+^ oscillations by PC2, channel-containing liposomes were flattened and scanned in a system we recently developed to simultaneously assess single channel currents and topological features of the ion channels (23). The oscillatory phenomenon occurred when PC2_iv_ clustering of various channel complexes was observed (Fig. 2b Bottom).

### Current oscillations of reconstituted PC2_iv_ in the presence of Ca^2+^ gradient and other ions

To determine whether the PC2_iv_ current oscillations were caused by Ca^2+^ transport, saline solutions in the reconstitution chamber were substituted with other salts. We first replaced the *cis* solution with 150 mM KCl. PC2_iv_ was reconstituted using the 3:7 POPC:POPE lipid mixture in this new biionic gradient condition (*cis* 150 mM KCl, 10 μM CaCl_2_ and *trans* 10 mM CaCl_2_). No oscillations were observed despite the fact that channel activity was present under these conditions. Subsequently, 200 μl of the *trans* solution was replaced by 200 μl of 1.0 M CaCl_2_, obtaining a final *trans* concentration of 210 mM CaCl_2_. With the increase of *trans* Ca^2+^, oscillations were again observed (*n* = 10/36, Fig. 3a Top). However, oscillations were different in shape from those observed in the presence of a Ca^2+^ gradient alone (see Fig. 2 for comparison). Because the oscillatory phenomenon was observed even in the absence of high *cis* Ca^2+^, we further evaluated if replacement of the *trans* solution (10 mM CaCl_2_) affected the appearance of the oscillations. PC2_iv_ channels were placed on the 3:7 lipid mixture under symmetrical 150 mM KCl and 10 μM CaCl_2_ Conditions. Although channel activity was observed, no ionic oscillations were elicited (data not shown). Subsequently, KCl was replaced by CaCl_2_ to the *trans* solution, obtaining a final *trans* concentration of 120 mM KCl and 200 mM CaCl_2_. With the increase in *trans* Ca^2+^, current oscillations were again observed (*n* = 4/14, Fig. 3a Bottom). Therefore, the amplitude, frequency and shape of the current oscillations through PC2_iv_ depended on the imposed ionic gradients, the applied holding potential, and the presence of high Ca^2+^ in the hemi-chambers.

### Kinetic model of PC2 channel simulated in QuB

From the results of the oscillatory currents elicited by reconstituted PC2 in the various preparations, a kinetic model was sought that could explain the oscillatory behavior of PC2_iv_. Our previous studies demonstrated that both PC2_hst_ and PC2_iv_ display four subconductance states of equal amplitude in a K^+^ gradient (150 mM *cis* and 15 mM *trans*), consistent with intrinsic characteristics of the ion channel (4, 5). Based on these findings, we designed a kinetic model of the PC2 channel in the QuB Classic suite (Open Ware, New York State Univ, Buffalo, NY). This software to design ion channel behavior is based on Markov processes, where the transitions between different substates of the channel are probabilities, such that changes per unit time of the state’s probability will only depend on the event in progress and not on past events (no memory). The PC2 kinetic model consisted of five states, a closed (C) state, and four conductive (open, O) substates O_1_ - O_4_ (Fig. 4a) where each of the states is characterized by increasing amplitude, and each transition by its speed constants (see Table 1). The simulations were run with *n* = 1 (Fig. 4b), and the simulation time was 9 seconds. This kinetic model of PC2 was the one that more closely approximated the previously reported data from our laboratory (Fig. 4b, see Fig. 1, in 5), and reproduced the mean closed time and the four sub-conductance states of the PC2_hST_ channel.

**Table 1.** Kinetic Parameters for the five substate PC2 model

| Substate | Current, pA | SD, pA | Transition constant to previous level, s<sup>-1</sup> | Transition constant to next level, s<sup>-1</sup> |
| --- | --- | --- | --- | --- |
| **C** | 0 | 0.01 | - | 1000 |
| **O<sub>1</sub>** | 1 | 0.05 | 500 | 1000 |
| **O<sub>2</sub>** | 2 | 0.05 | 500 | 1000 |
| **O<sub>3</sub>** | 3 | 0.05 | 250 | 1000 |
| **O<sub>4</sub>** | 4 | 0.05 | 100 | - |

**Fig. 4:**
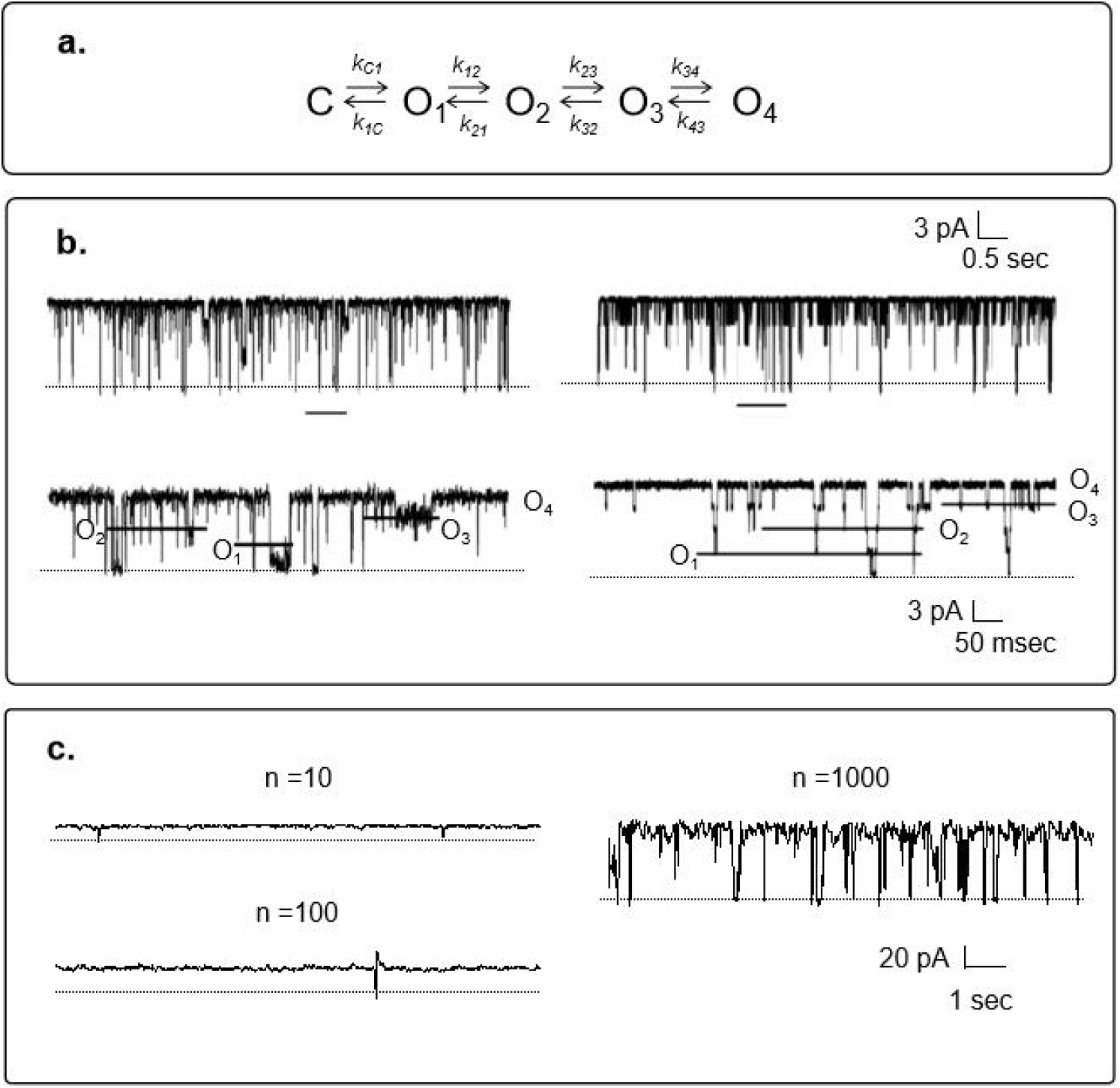
QuB simulation of PC2 channel behavior. a. Kinetic model used in the QuB simulation of PC2-mediated K^+^ currents. b. Left. PC2_hst_ single channel currents in a K+ gradient (150 mM cis KCl, 15 mM trans) and a holding potential of 60 mV as originally reported (5). Tracing below is an expanded region indicated at the horizontal line, where the four substates (O1-O4) of the channel are indicated by the horizontal lines. Right. Simulated kinetic model of PC2 in QuB. Horizontal lines indicate the different subconductance levels (O_1_-O_4_) of the simulated tracing. Dotted lines indicate zero current level. c. Simulation with n = 10, 100 and 1000 channel units. Dotted line indicates zero current level.

### Simulation of PC2 channel oscillations with QuB

To determine whether the activity of the PC2_QuB_ channels (PC2 reconstructed by QuB) kinetically reproduced the oscillations observed under Ca^2+^ gradients and biionic conditions (K^+^ and Ca^2+^), several channels were idealized with the same parameters. From 2 to 100 identical channels were explored with the same Markovian characteristics. However, the simulation failed to reproduce the current oscillations, even by increasing channel number to *n* = 1000 (Fig. 4c). This simulated model responded to the expected stochastic model.

The experimentally-observed PC2-mediated Ca^2+^ current oscillations showed large amplitudes that depended on the imposed gradient. In all cases, the amplitude was greater than 50 pA, meaning that oscillations required the synchronous activity of several open channels. During oscillations it was not possible to visualize the four conductive states of the channel. All the channels must be completely opened at the *g*_*max*_ level, and closed in a synchronized manner (see Figs. 2 & 3). The previously proposed stochastic model, which reproduced the PC2 single channel currents in a K^+^ gradient, failed to reproduce the multiple channel oscillations, therefore a mechanism was further sought that could better approximate the experimental data. Thus, another kinetic model of the channel was designed, which was called a PC2_iv_ oscillatory model (Fig. 5a), which consisted of a closed state and three open states of equal amplitude (see Table 2). The rate constants for each transition state were modified such that when changing the residence time, the behavior of the channel simulated a cooperative phenomenon (Table 2). Thus, values for the transition constants were chosen such that they reproduced a rapid opening of the channel to its *g*_*max*_ level, and then also quickly fell to the closed state. These channel transitions generated an oscillatory behavior.

**Table 2.** Transition constants of the four substate PC2 kinetic model

| Substate | Current, pA | SD, pA | Transition constant to previous level, sec<sup>-1</sup> | Transition constant to next level, sec<sup>-1</sup> |
| --- | --- | --- | --- | --- |
| C | 0 | 0.10 | - | 10000 |
| O<sub>1</sub> | 10 | 0.15 | 1 | 10000 |
| O<sub>2</sub> | 10 | 0.15 | 100 | 1 |
| O<sub>3</sub> | 10 | 0.15 | 1000 | - |

**Fig. 5:**
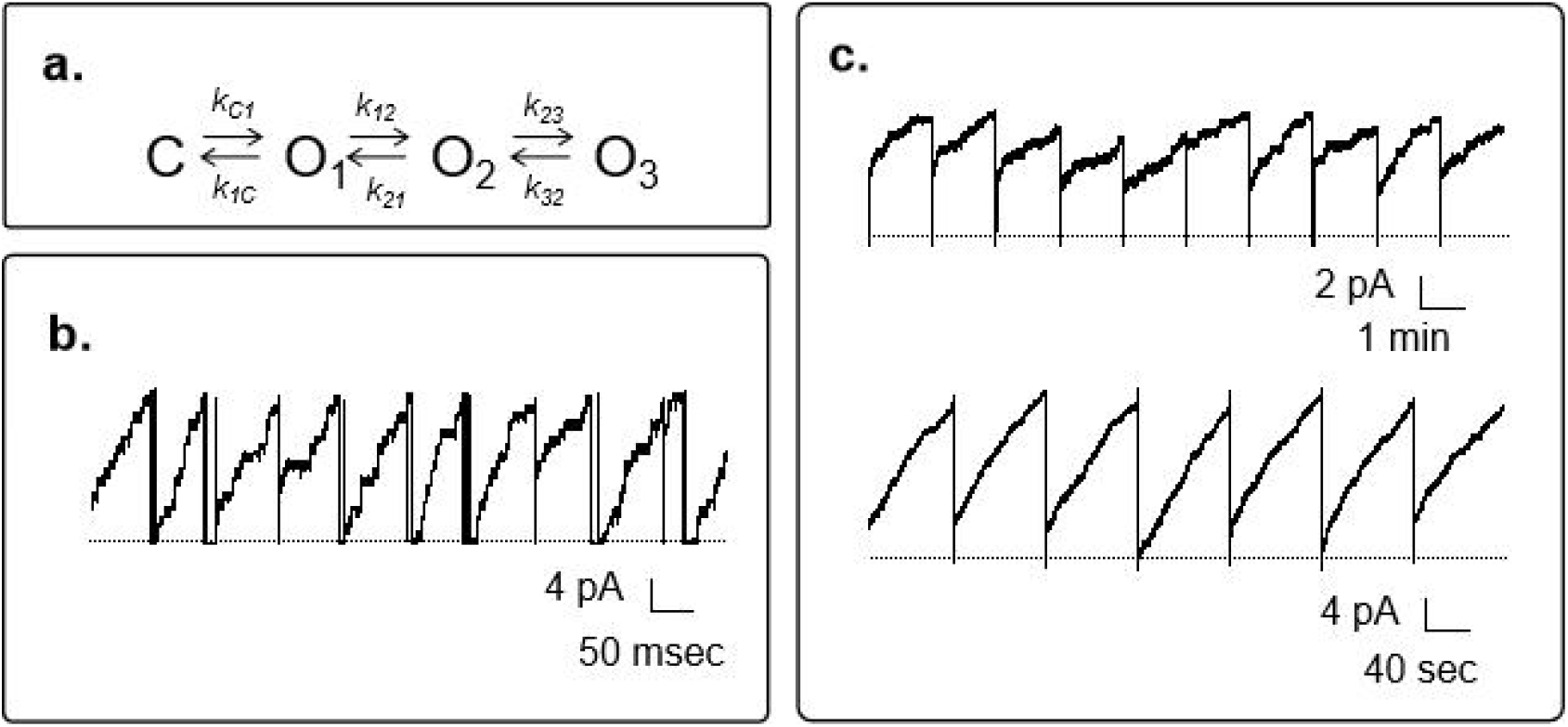
QuB simulation model of PC2-mediated current oscillations. a. Kinetic model of PC2 that supports current oscillations. b. Representative current oscillations simulated with 100 channels with 10 pA current amplitude. c. Simulation with 100 and 1000 channels (Top and Bottom, respectively) of the same kinetic model, with PC2 channels whose open state amplitude is 1 pA.

A representative plot of current oscillations in symmetrical K^+^ was observed where the kinetic model partially reproduced the periodic behavior of the PC2 channel when the simulation ran with *n* = 100 channels (Fig. 5b). Simulations of the PC2 channel with the same kinetic model, but a modified conductance to a tenth of the original conductance, i.e. 1 pA maximum amplitude, also managed to reproduce the oscillatory behavior when running simulations with 100 and 1000 channels (Fig. 5c). Thus, the oscillatory behavior of the PC2 channel currents largely depends on the kinetic changes of the channel, and not to changes in its conductance. Amplification of PC2 Ca^2+^ currents in a Ca^2+^ gradient and the QuB simulated oscillations shows the staircase behavior in both cases (Fig. 6 a & b)

**Fig. 6:**
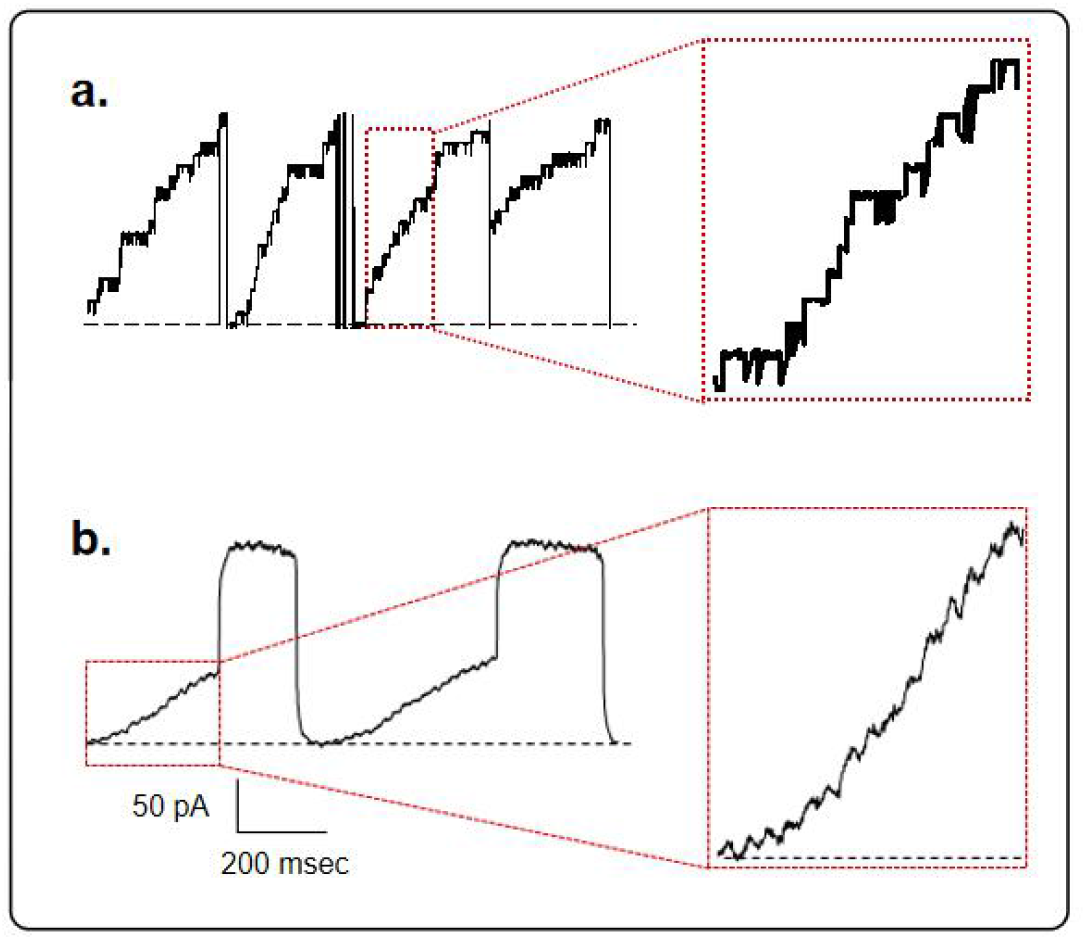
Staircase amplification of PC2 Ca^2+^ currents. a. Left. QuB simulated oscillations with a 100, 1 pA channels. Right. Amplification of PC2_ptv_ Ca^2+^ currents. b. Left. Representative tracing of PC2 current oscillations in a Ca^2+^ gradient, and its corresponding amplification (Right). Holding potential 117 mV. The inset shows unitary jumps.

## Discussion

The regulation by Ca^2+^ of Ca^2+^-permeable ion channels represents an important mechanism in the control of all cellular function. The carboxyl terminus of the epithelial Ca^2+^ channel ECaC1, for example, is involved in Ca^2+^ dependent inactivation (30). PC2, a member of the TRP channel superfamily, is a high conductance, Ca^2+^-permeable non-selective cation channel, whose dysfunction causes ADPKD. The study of Ca^2+^ transport and channel regulation by Ca^2+^ through this and other channels expressed in non-excitable cells, is of relevance, because as a whole, it represents one of the initial steps in signaling mechanisms of the primary cilium and other cellular compartments (31).

The present study evaluated Ca^2+^ transport by PC2 at high Ca^2+^ concentrations. Reconstituted PC2_iv_ in a Ca^2+^ gradient elicited spontaneous Ca^2+^ currents in 64% of the cases (*n* = 105/360). PC2 channel reconstitution in a BLM system provided us the means to determine that the presence of different phospholipid compositions, namely, POPC: POPE, 3:7 and 7:3, in the membrane, affected the PC2 mediated Ca^2+^ currents in the isolated protein (PC2_iv_). Significant differences were observed in the maximum Ca^2+^ conductance, which was almost twice as large in the 7:3 POPC: POPE ratio, as compared to the reverse 3:7 ratio; thus, suggesting an important effect of the lipidic environment on channel conductance. Interestingly, the estimated V_rev_ in both conditions was unaffected by the lipidic environment, and consistent with the same high cationic perm-selectivity. The truncated PC2_R742X_ channel, also displayed similar Ca^2+^channel activity, suggesting that the absence of the carboxy-terminal tail of the channel does not affect Ca^2+^ transport by PC2. However, PC2_R742X_ channel was more difficult to reconstitute in a BLM and obtain Ca^2+^ currents, probably because high Ca^2+^ affect the stability of the interaction between the subunits that form the channel pore. Moreover, no oscillations were observed with this sample.

A surprising observation while studying PC2_iv_ Ca^2+^ transport in Ca^2+^ gradients was the presence of ionic current oscillations. This oscillatory behavior was observed in 21% (*n* = 29/139) of the experiments where PC2_iv_ was reconstituted in the presence of the 3:7 POPC: POPE lipid mixture, but not the 7:3 lipid mixtures. Neither were oscillations observed with truncated PC2_R742X_, suggesting that the carboxy terminal end of the protein might play a role in the phenomenon, most likely by contributing to the expected ion channel assemblies. PC2 channel clustering on the oscillatory ion channel current phenomenon was further confirmed by current recordings of AFM scanned BLMs, with the technique we recently reported (23). The amplitude and intrinsic frequency of the current oscillations depended on various parameters, including the applied holding potential, the imposed ionic gradient, and the presence of high Ca^2+^. This interesting phenomenon, likely generated by the clustering of a single ion channel species in the absence of extrinsic regulatory proteins, requires a feedback mechanism, as expected from other Ca^2+^ chemical oscillators, and in this case, actually depended upon the transport kinetics of the channel that is controlled by the same transported Ca^2+^. This hypothesis was confirmed by simulations of multistep PC2 channel behavior in the QuB program, where we observed that the simple stochastic behavior of PC2_iv_ would respond as a quasi-deterministic phenomenon, as observed in stochastic resonance (32).

The data in this study provides novel evidence, confirming the Ca^2+^-transporting capabilities of PC2, further supporting the contention that the channel could be self-regulated by means of feedback mechanisms, which are independent of external regulatory proteins, as previously observed in our laboratory (17). This oscillatory behavior, previously unknown for single channel species, would likely depend on the presence of Ca^2+^ interaction sites as have been postulated for the carboxy terminus of the channel protein (18). The same region in the PC2-like, polycystin-L (33), however, seems to convey the same effect. Our results refute the apparent absence of regulatory sites in the channel protein, which is unapparent at low Ca^2+^concentrations. The oscillations in the Ca^2+^ currents of the isolated protein (PC2_iv_) would require interaction sites, as a feedback mechanism that generates the oscillations, where the kinetics of the channel would be controlled by the transported ion. This may involve a coupling of the ion to intracellular sites of the same channel or, potentially, “structural” phospholipids, or charges that control the probability of the open state. Since the oscillatory electrical responses in “non-excitable” cellular models would be associated with oscillations of intracellular Ca^2+^ (13, 14), the data in this study, suggest a minimalist model where the voltage sensitivity of PC2 would be conferred by Ca^2+^-dependent conformational changes. From these findings, we have been able to confirm the importance of Ca^2+^ permeability by PC2, and its regulation by the ion, in a behavior that is comparatively similar to voltage-activated Ca^2+^ channels. Future studies will be required to further explore the possible feedback mechanisms that help determine whether the presence of the oscillatory phenomenon observed with the isolated PC2_iv_ protein is also possible in different cellular environments.

## Author contributions

IFV carried out all experimental procedures. IFV and MdRC conducted the analysis of the experimental data and prepared Figures and Tables. HFC and MdRC designed all the experiments and wrote the main manuscript. All authors reviewed the manuscript.

## Acknowledgements

This study was funded by Ministerio de Ciencia, Tecnología e Innovación, Argentina, FONCyT, PICT 2012 N°1559 (HC).

